# The role of hepatitis B virus surface protein in inducing Sertoli cell ferroptosis

**DOI:** 10.1101/2022.03.24.485732

**Authors:** Chengshuang Pan, Kong xiangbin, Wu zhigang, Qianjin Fei

**Author notes:** Address correspondence to **Qianjin Fei**, j. No. 96 Fuxue Xiang, Lucheng District, Wenzhou, Zhejiang, 325000, China.

## Abstract

Hepatitis B virus infection could result in male infertility by inhibiting sperm function and viability. Sertoli cell death contributes to spermatogenesis impairment, which is associated with sperm defects and dysfunction. Ferroptosis-mediated cell death of Sertoli cells was found to contribute to spermatogenesis disorder and poor sperm quality. However, the effects of hepatitis B virus infection on ferroptosis of Sertoli cells remain to be elucidated. Human Sertoli cells were cultured in vitro with 25, 50, and 100 mg/mL of hepatitis B virus surface protein for 48 hours. Cell viability was measured with CCK-8. Levels of glutathione, malondialdehyde, iron, and m6A in human Sertoli cells were determined. Lipid peroxidation was assessed using C11-BODIPY. Luminescence analysis was performed to detect the binding of METTL3 and 3¢-UTR of TRIM37 containing the m6A motifs. Immunoprecipitation was applied to determine the relationship between TRIM37 and GPX4. qPCR and immunoblotting were performed to measure mRNA and protein levels. Hepatitis B virus surface protein exposure significantly increased TRIM37 expression, malondialdehyde level, and ferroptosis, and decreased cell viability and glutathione level of human Sertoli cells. TRIM37 silencing inhibits the effect of HBs exposure-regulated cell viability and ferroptosis in human Sertoli cells. TRIM37 inhibits GPX4 expression through ubiquitination. GPX4 overexpression inhibits the effect of TRIM37 on cell viability and ferroptosis in human Sertoli cells.

Administration of ferroptosis inhibitor recovers the cell viability decreased by TRIM37. Mechanism study showed HBs increases the level of TRIM37 3’-UTR m6A by promoting the expression of METTL3, and the binding of m6A reader IGF2BP2 and TRIM37 3’-UTR promotes the stability of TRIM37 mRNA.HBs inhibit Sertoli cell viability by promoting ferroptosis of Sertoli cells through TRIM37-mediated ubiquitination of GPX4. The findings highlight the importance of TRIM37/GPX4 signaling in the ferroptosis of Sertoli cells.

## Introduction

Infertility has been a concern for ages and is also a significant clinical problem that affects about 10% of couples worldwide. “Malefactor” infertility accounts for half of the cases(1). Deterioration of sperm quality increases the risks of embryo developmental disorders (2). Reasons for male subfertility could be related to factors that impair spermatogenesis and lead to poor sperm parameters. Semen analysis and functional analysis of sperm are the best diagnostics(3).

Many factors may involve in male infertility, including Hepatitis B virus (HBV) infection. HBV is a hepatotropic virus that can cause a chronic infection. A study showed that 3.5% of the population is HBV-infected (4). China has the highest HBV infection rate (9%)(5). HBV surface antigen (HBsAg) expression can trigger T cell dysfunction and favor detrimental immunopathology(6). Since HBV could pass through the blood-testis barrier and enter the testis, it may pose a risk to spermatogenesis. It has been shown that HBV up-regulated seminal plasma malondialdehyde (MDA), enhanced oxidative stress, elevated IL-17 and IL-18, and resulted in male infertility (7). Moreover, HBV infection reduced sperm function and viability (5). However, the mechanism underlying the adverse effect of HBV on spermatogenesis and sperm quality remains to be elucidated.

Sertoli cells, which are the only somatic cells located inside the seminiferous epithelium (8), are major supportive germ cells that regulate spermatogenic cells via several paracrine factors (9). Sertoli cell death contributes to spermatogenesis impairment that can cause sperm defects, and result in male infertility (10).

Developing a better understanding of the signaling pathway that induces Sertoli cell death is important for the development of an effective strategy to prevent germ cell loss and infertility.

Ferroptosis is non-apoptotic cell death that can be found in various diseases, including ischemia-reperfusion injury(10). For example, ferroptosis plays an important role in cardiomyopathy and heart failure (11). Hepatic ferroptosis has also been shown to trigger inflammation in nonalcoholic steatohepatitis (12). Bromfield et al. also showed oxidative stress promotes ferroptosis in spermatids (13). More importantly, oxygen/glucose deprivation and re-oxygenation cause ferroptosis in Sertoli cells to contribute to male infertility (10). Knowing the underlying mechanism of Sertoli cell death is important for preventing spermatogenic arrest(10).

Tripartite motif (TRIM)-containing proteins are a family of approximately 70 proteins that have an N-terminal RING finger, one or two B-boxes, and a coiled-coil (CC) domain (14). The RING finger structure of TRIM is closely related to E3 ubiquitin ligases (E3s) activity(15). Research suggested that TRIM proteins are closely related to the pathogenesis of various diseases. Hou et al. have reported that loss of TRIM21 ameliorates cardiotoxicity via inhibiting ferroptosis (16). Many TRIM proteins have been shown to regulate different cellular biological processes including ferroptosis via regulating p53 levels and activity (17). TRIM16L, TRIM37, TRIM40, TRIM56, and TRIM59 have been shown to regulate HBV viral protein expression (18). Studies also showed that TRIM22 contributes to the clearance of HBV (19). HBV X protein (HBx) was demonstrated to upregulate TRIM7 expression, leading to the highly proliferative characteristics of hepatocellular carcinoma cells (20).

Regardless of the great advances made in the field of HBV and ferroptosis, the effect of HBV on ferroptosis of Sertoli cells remains to be elucidated. Therefore, we explored the role of HBV in the ferroptosis of Sertoli cells.

## Results

### HBs exposure decreased cell viability and increased ferroptosis of Sertoli cells

To study the effects of HBs on Sertoli cells, human Sertoli cells were treated by HBs at different concentrations (25, 50, and 100 mg/mL). Results showed that HBs treatment inhibited the viability of human Sertoli cells (Figure 1A), enhanced cellular iron concentration (Figure 1B), and promoted lipid peroxidation, indicated by elevated fluorescence of C11-BODIPY and MDA level (Figures 1C-1D) in a dose-dependent manner. In contrast, HBs treatment dose-dependently decreased GSH level (Figure 1E) and inhibited the expression of GPX4 (Figure 1F). Together, these findings show that HBs exposure decreased cell viability with increasing ferroptosis of Sertoli cells.

**Figure 1.**
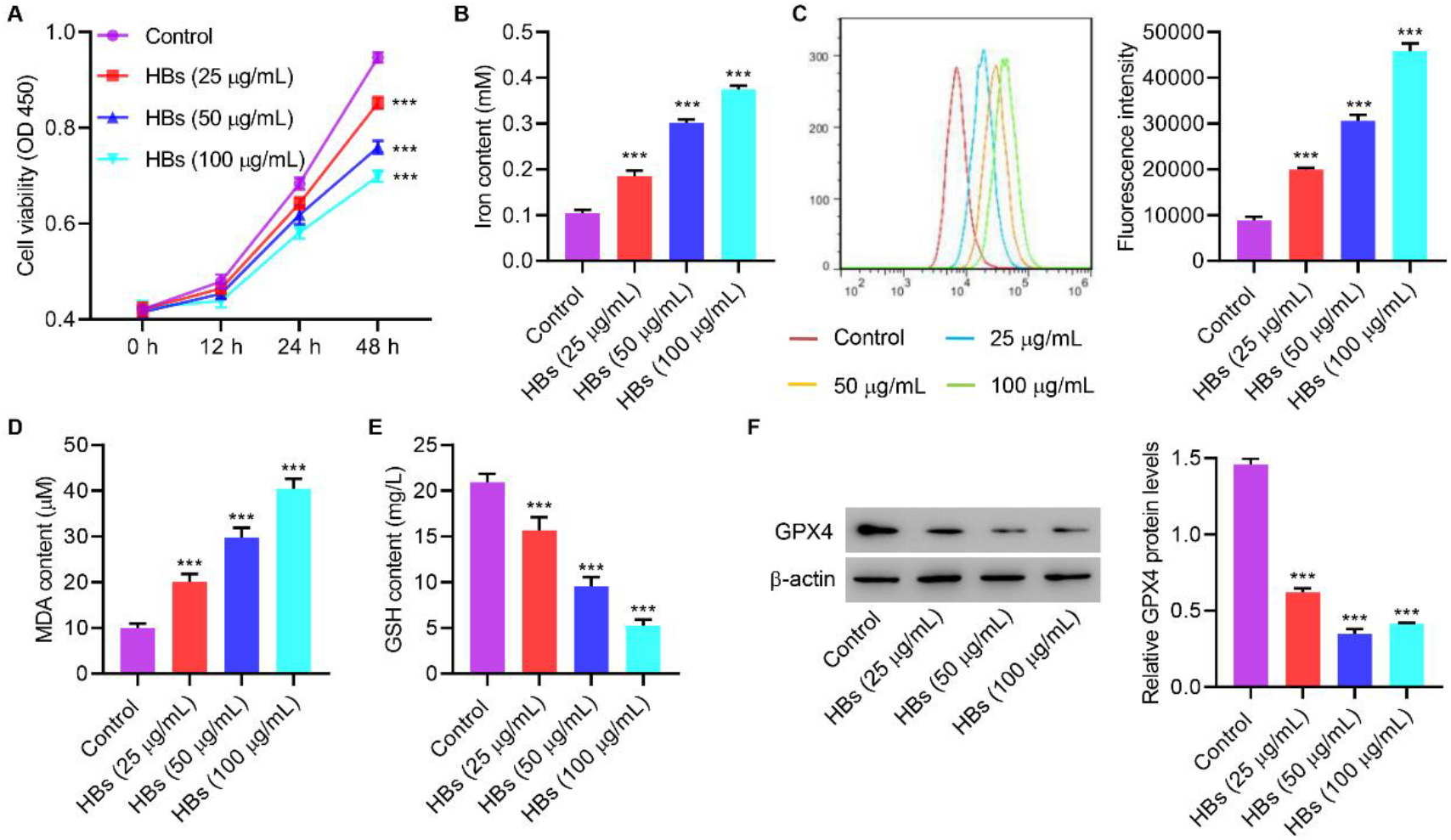
Effects of HBs exposure on viability and ferroptosis of Sertoli cells. Cells were provided HBs (25, 50, and 100 μg/mL), and the (A) cell viability, (B) iron content, (C) lipid peroxidation, (D) MDA, (E) GSH, and (F) GPX4 expression were measured. \*\*\**P*<0.001 compared with control.

### TRIM37 silencing inhibits effects of HBs exposure regulated cell viability and ferroptosis in human Sertoli cells

To evaluate the role of TRIM in HBs exposure-regulated cell viability and ferroptosis of Sertoli cells, we first measured TRIM expression. Results showed that HBs exposure dose-dependently increased expression of TRIM37 compared to other TRIMs (Figure 2A). Next, TRIM37 was successfully silenced (Figures 2B-2C).

**Figure 2.**
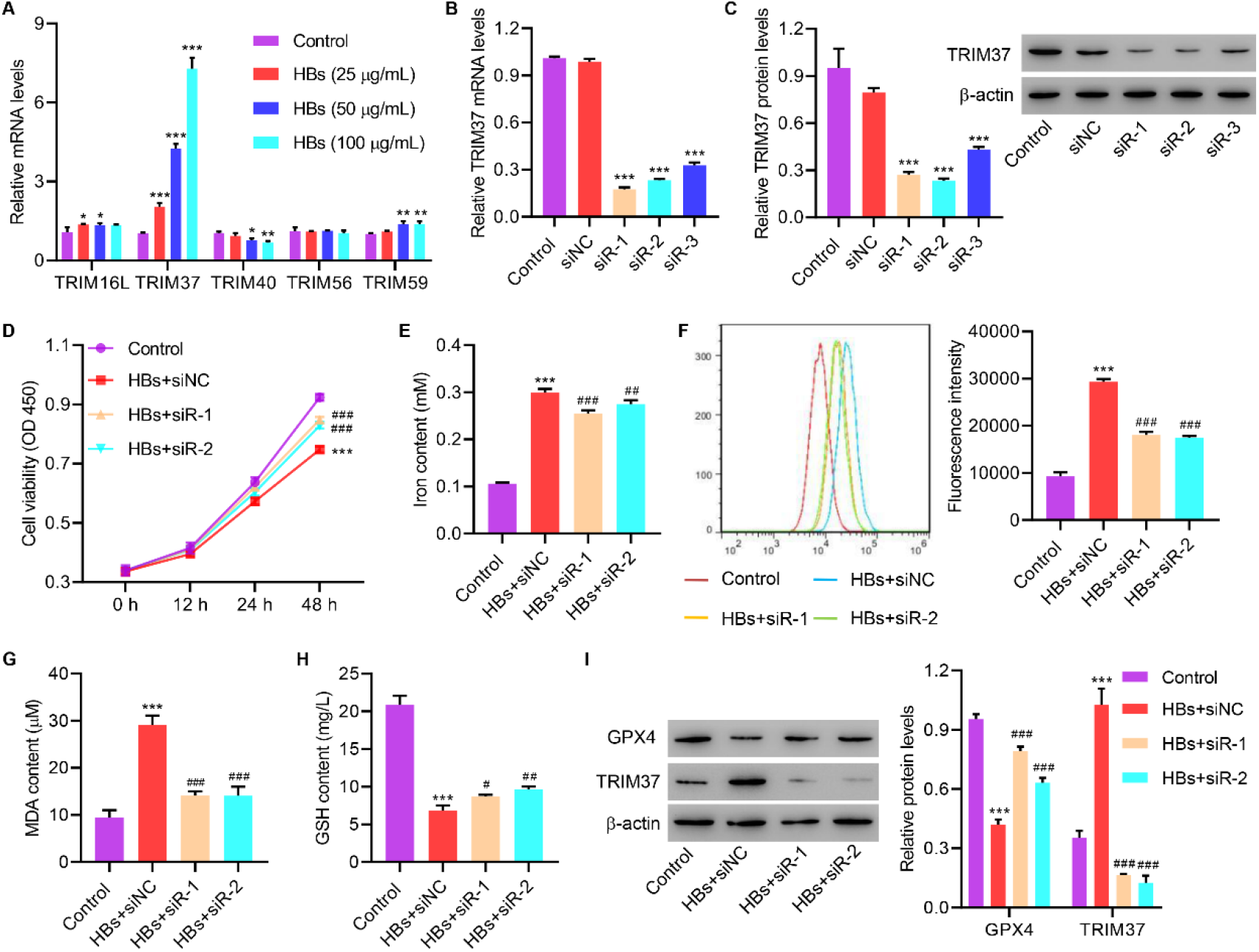
TRIM37 silencing inhibits HBs’ effects on viability and ferroptosis of Sertoli cells. (A) Cells were treated with HBs (25, 50, and 100 μg/mL), and the expression of TRIM16L, TRIM37, TRIM40, TRIM56 and TRIM59 was measured. Cells were transfected with TRIM37 siRNA, and TRIM37 expression was measured (B, C). Cells were treated with HBs (50 μg/mL) and transfected with TRIM37 siRNA, and the (D) cell viability, (E) iron content, (F) lipid peroxidation, (G) MDA, (H) GSH, and (I) expression of GPX4 and TRIM37 were measured. \*\*\**P* < 0.001 vs. ctrl. ^#^*P* < 0.05, ^##^*P* < 0.01, ^###^*P* < 0.001 vs. HBs+siNC.

Silencing TRIM37 significantly ameliorated HBs exposure-decreased viability (Figure 2D) and HBs-exposure-enhanced cellular iron concentration (Figure 2E) and lipid peroxidation (Figures 2F-2G). Silencing TRIM37 also significantly abolished HBs-exposure-decreased GSH level (Figure 2H) and GPX4 expression (Figure 2I).

The results suggest that TRIM37 silencing inhibits the effect of HBs exposure-regulated cell viability and ferroptosis of human Sertoli cells.

### TRIM37 inhibits GPX4 expression through ubiquitination

To further explore the relationship between TRIM37 and GPX4, TRIM37 was successfully overexpressed in human Sertoli cells (Figures 3A-3B). Overexpression of TRIM37 significantly decreased the protein level of GPX4, while silencing TRIM37 significantly increased the level of GPX4. In contrast, neither silencing nor overexpressing TRIM37 affected GPX4 at the mRNA level (Figures 3C-3D). Administration of proteasome inhibitor, MG132, significantly abolished TRIM37 overexpression-caused down-regulation of GPX4 (Figure 3E). Immunoprecipitation results showed that TRIM37 interacted with GPX4 (Figure 3F) and overexpressing TRIM37 significantly increased ubiquitination of GPX4 (Figure 3G). These results demonstrate that TRIM37 inhibits GPX4 expression through ubiquitination.

**Figure 3.**
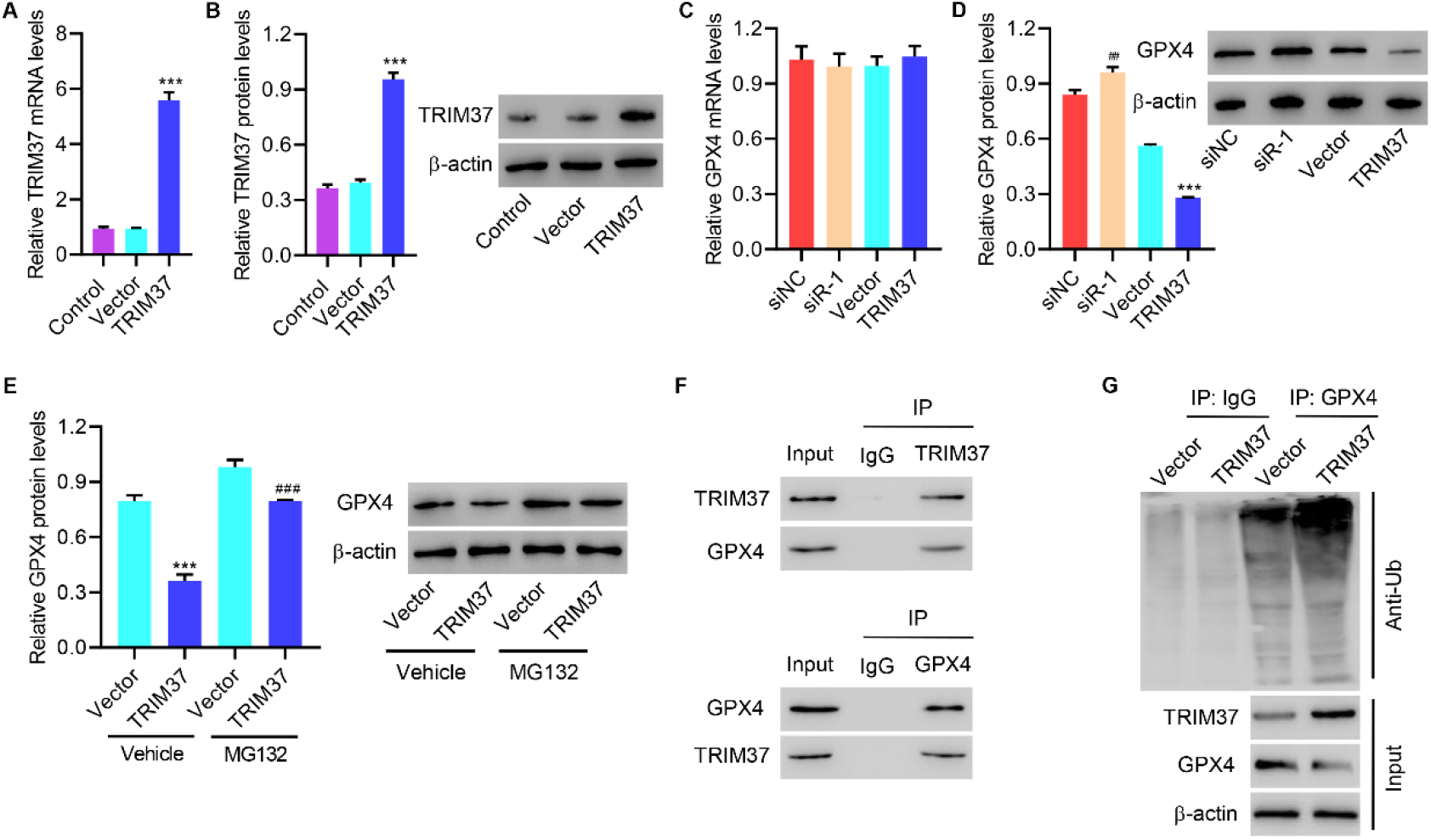
TRIM37 inhibits GPX4 expression through ubiquitination of GPX4. (A, B) Cells were transfected with TRIM37 expression plasmid, and TRIM37 expression was measured (C-E). Cells were transfected with TRIM37 siRNA or expression plasmid with 10 μM MG132 or its vehicle treatment, and TRIM37 expression was measured. (F) IP was performed with a control IgG, anti-TRIM37, or anti-GPX4 antibody, followed by incubation with indicated antibodies. (G) TRIM37-overexpressing human Sertoli cells immuneprecipitated with GPX4 or IgG antibody for evaluating ubiquitination. \*\*\**P*<0.001 compared with vector or vector+vehicle. ^##^*P*<0.01, ^###^*P*<0.001 compared with siNC or TRIM37+vehicle.

### GPX4 overexpression inhibits the effect of TRIM37 on the cell viability and ferroptosis in Sertoli cells

To further clarify the roles of GPX4, we successfully overexpressed GPX4 in TRIM37-overexpressing human Sertoli cells (Figures 4A-4B). Overexpressing GPX4 abolished TRIM37 overexpressing-suppressed cell viability (Figure 4C), enhanced lipid peroxidation (Figures 4D-4E), and suppressed GPX4 expression (Figure 4F).

**Figure 4.**
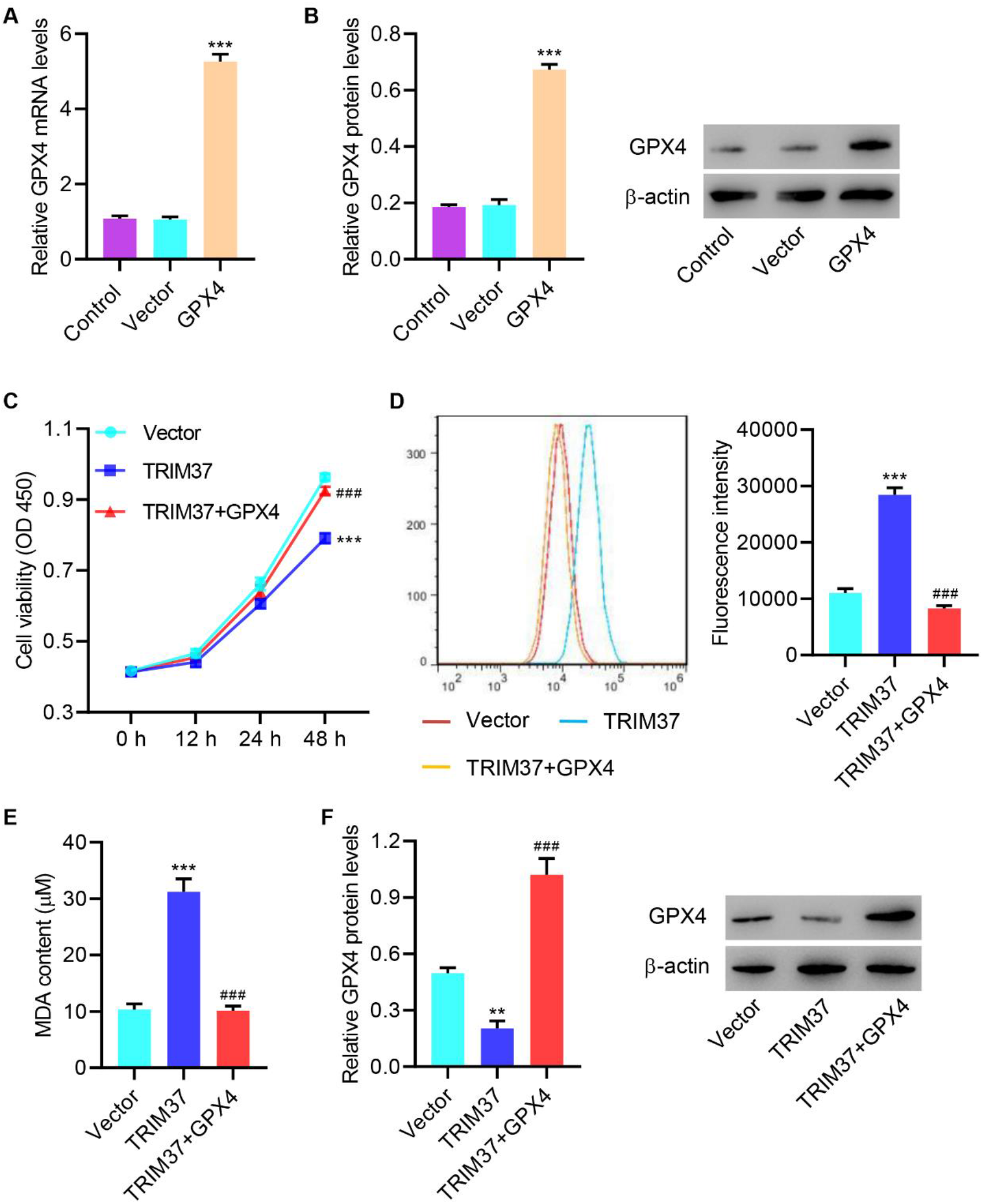
GPX4 overexpression inhibits the effect of TRIM37 on the cell viability and ferroptosis. (A, B) Cells were transfected with GPX4 expression plasmid, and GPX4 expression was measured. Cells were transfected with TRIM37 expression and GPX4 expression plasmid, and the (C) cell viability, (D) lipid peroxidation, (E) MDA, and (F) GPX4 expression were measured. \*\**P* < 0.01, \*\*\**P* < 0.001 vs. vector. ^###^*P* < 0.001 vs. TRIM37.

Together, data indicates that GPX4 overexpression suppresses the effects of TRIM37 on cell viability and ferroptosis in human Sertoli cells.

### Inhibition of ferroptosis recovers the cell viability decreased by TRIM37

To further explore the roles of ferroptosis in the viability of Sertoli cells, we introduced DFO, a ferroptosis inhibitor. Administration of DFO abolished TRIM37 overexpression-decreased Sertoli cell viability (Figure 5A) and significantly decreased TRIM37-enhanced lipid peroxidation (Figure 5B). These findings suggest that inhibition of ferroptosis recovers the cell viability decreased by TRIM37 in human Sertoli cells.

**Figure 5.**
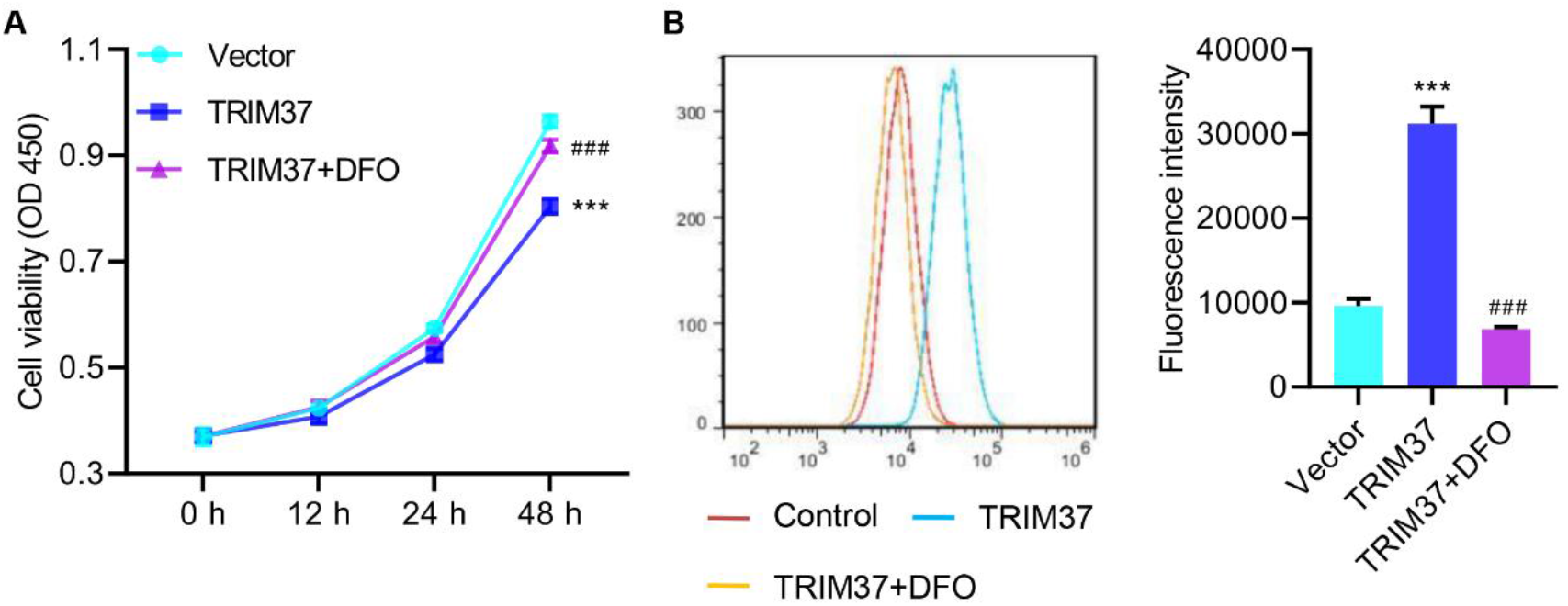
Inhibition of ferroptosis recovers the cell viability decreased by TRIM37 in human Sertoli cells. Human Sertoli cells were transfected with TRIM37 expression and treated with 100 μM iron chelators (deferoxamine; DFO), and the (A) cell viability and (B) lipid peroxidation were measured. \*\*\**P*<0.001 vs. vector. ^###^*P*<0.001 vs. TRIM37.

### METTL3-mediated m6A modification caused the upregulation of TRIM37 via IGF2BP2 in HBs-induced human Sertoli cells

Bioinformatic analysis revealed that METTL3 might mediate the methylation of TRIM37 mRNA, so we checked the effect of HBs on m6A methylation level. Results showed that HBs exposure dose-dependently increased m6A methylation level (Figure 6A) and TRIM37 at both mRNA and protein levels (Figures 6B-6C). Next, the m6A methylation level of TRIM37 was determined by MeRIP-qPCR assays and the result showed that HBs exposure significantly increased m6A methylation of TRIM37 (Figure 6D). Then, human Sertoli cells were treated with HBs (50 mg/mL) and transfected with METTL3 siRNA or expression plasmid (Figure S1A-1B).

**Figure 6.**
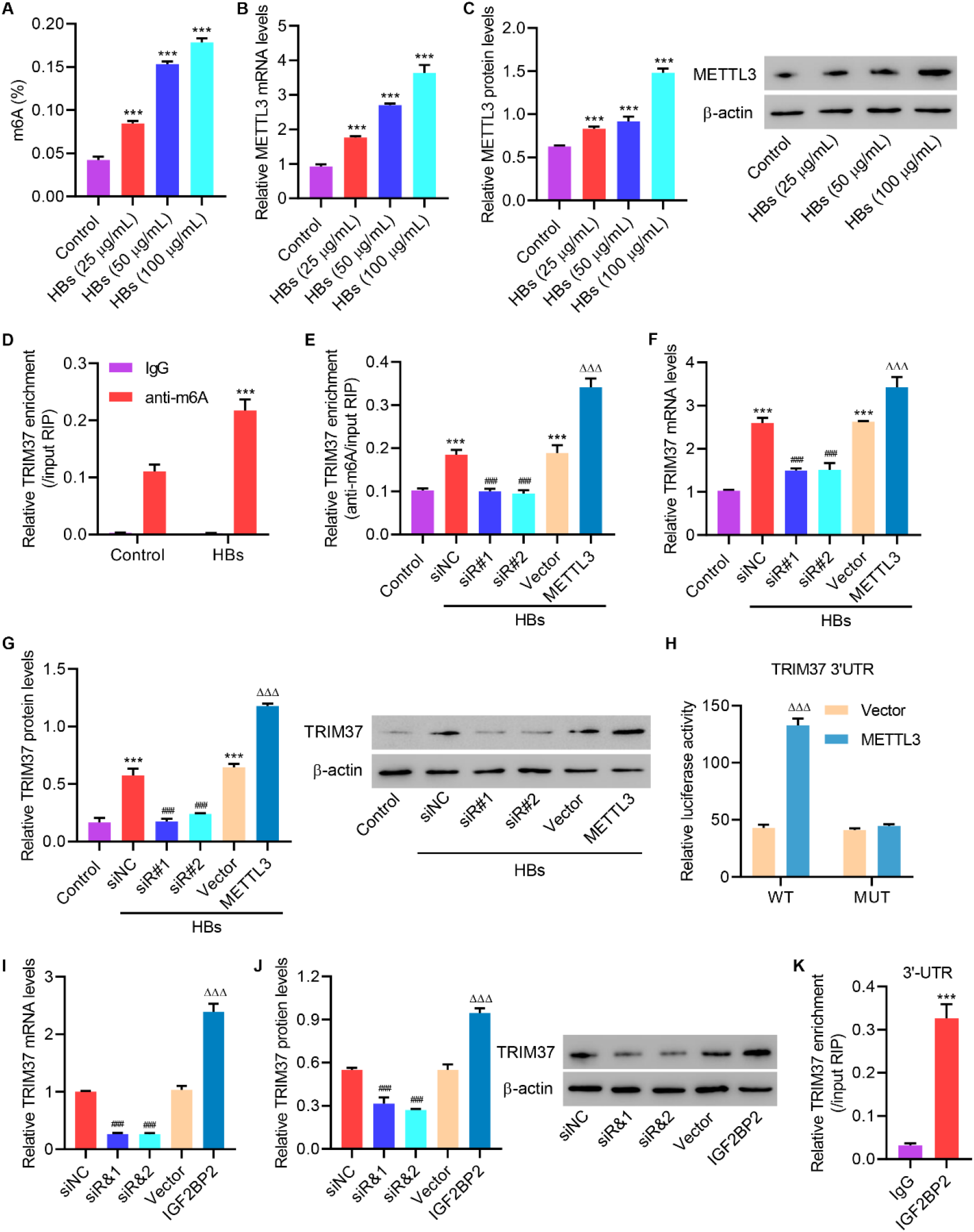
METTL3-mediated m6A modification causes the upregulation of TRIM37 via IGF2BP2 in HBs-induced Sertoli cells. (A) Cells were treated with HBs (25, 50, and 100 μg/mL), and the m6A methylation level was measured. (B, C) Cells were treated with HBs (25, 50, and 100 μg/mL), and METTL3 expression was measured. (D) The m6A methylation level of TRIM37 was determined by MeRIP-qPCR assays. Human Sertoli cells were treated with HBs (50 μg/mL) and transfected with METTL3 siRNA or expression plasmid, and the (E) m6A methylation level of TRIM37 and (F, G) expression of TRIM37 were measured. (H) WT or mutant m6A consensus sequences within TRIM37 3’-UTR were fused with a firefly luciferase reporter. Sertoli cells transfected with an empty vector or METTL3 expression plasmid were transfected with a luciferase reporter plasmid. Luciferase activity was measured. Cells were transfected with IGF2BP2 siRNA or expression plasmid, and the (I, J) expression of TRIM37 were measured. (K) Enrichment of TRIM37 3’-UTR following immunoprecipitation of IGF2BP2 from extracts of human Sertoli cells. \*\*\**P*<0.001 vs. control or IgG. ^###^*P*<0.001 vs. HBs+siNC or siNC. ^ΔΔΔ^*P*<0.001 vs. HBs+vector or vector.

Silencing METTL3 dramatically down-regulated m6A methylation level of TRIM37, while METTL3 overexpression sharply up-regulated m6A methylation level of TRIM37 (Figure 6E). Silencing METTL3 also significantly decreased TRIM37, while overexpressing METTL3 significantly increased TRIM37 (Figures 6F-6G). Next, WT or mutant m6A consensus sequences within TRIM37 3’-UTR were fused with a firefly luciferase reporter and used for transfection of human Sertoli cells with either an empty vector or METTL3 expression plasmid. Luciferase activity measurement showed that METTL3 overexpression significantly increased luciferase activity in WT TRIM37 3’-UTR reporter plasmids in Sertoli cells. However, METTL3 overexpression showed no significant effects on luciferase activity in mutated TRIM37 3’-UTR reporter plasmids (Figure 6H). Then, Sertoli cells were transfected with m6A reader, IGF2BP2, siRNA, or expression plasmid (Figure 7C-7D). Silencing IGF2BP2 significantly decreased TRIM37, while overexpressing IGF2BP2 significantly increased TRIM37 (Figures 6I-6J). The interaction between TRIM37 and IGF2BP2 was also confirmed (Figure 6K). All these findings demonstrated that HBs increase the level of TRIM37 3’-UTR m6A by promoting the expression of METTL3, and the binding of m6A reader IGF2BP2 and TRIM37 3’-UTR promotes the stability of TRIM37 mRNA.

**Figure 7.**
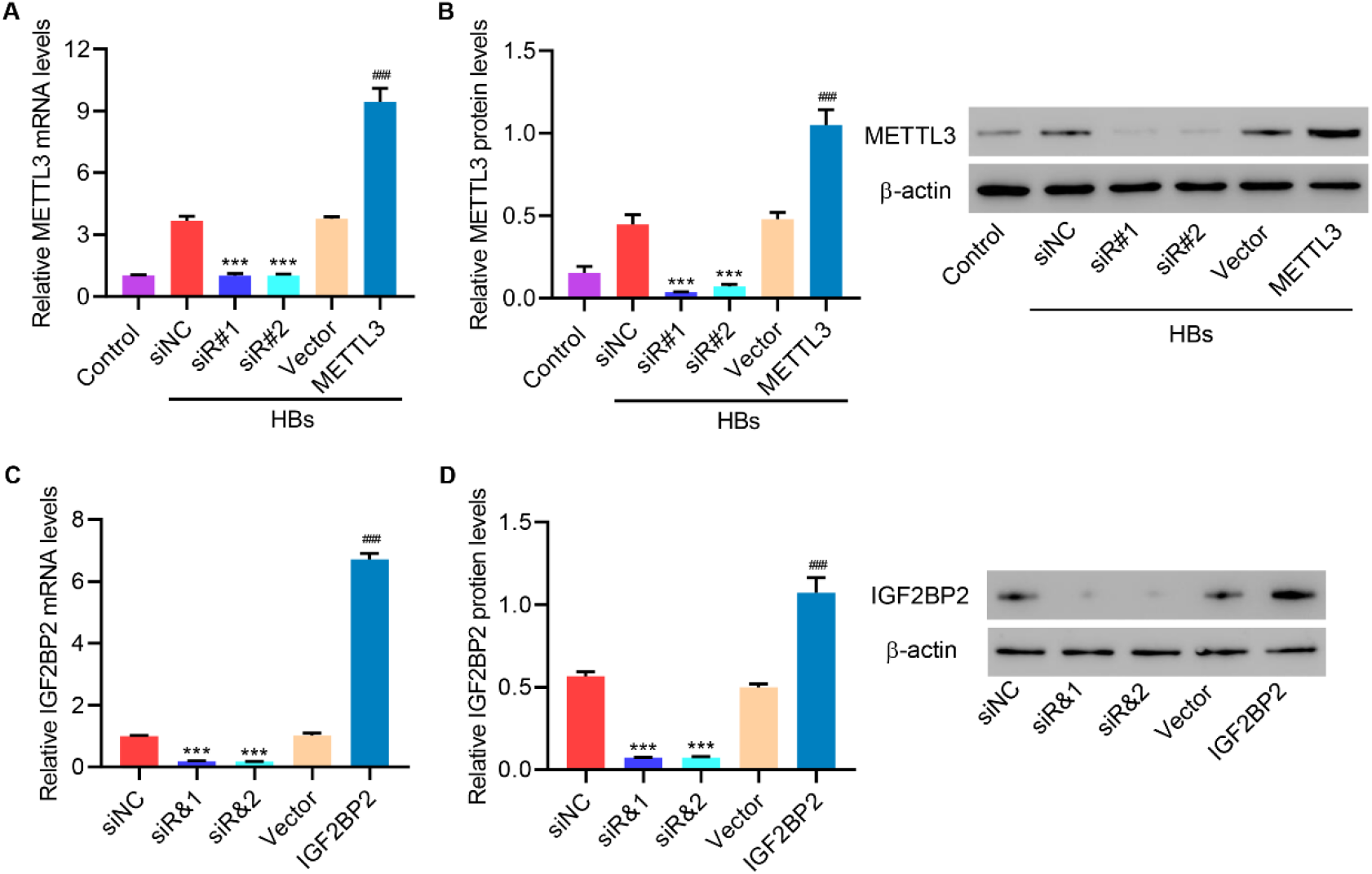
METTL3 and IGF2BP2 expression in Sertoli cells. (A, B) Cells were treated with HBs (50 μg/mL) and transfected with METTL3 siRNA or expression plasmid, and the expression of METTL3 was measured. (C, D) Cells were transfected with IGF2BP2 siRNA or expression plasmid, and the expression of IGF2BP2 was measured. \*\*\**P*<0.001 vs. HBs+siNC or siNC. ^###^*P*<0.001 vs. HBs+vector or vector.

## Discussion

In this study, we first analyzed the effects of HBs exposure on Sertoli cells and found that HBs exposure significantly decreased viability and increased ferroptosis of human Sertoli cells. HBs exposure also dose-dependently increased expression of TRIM37. TRIM37 silencing inhibits the effect of HBs exposure regulated cell viability and ferroptosis in human Sertoli cells. Our results also support that TRIM37 inhibits GPX4 expression through ubiquitination and GPX4 overexpression inhibits the effect of TRIM37 on the cell viability and ferroptosis in human Sertoli cells.

Mechanism study indicated that inhibition of ferroptosis recovers the cell viability decreased by TRIM37 in human Sertoli cells. Data also showed that HBs increase the level of TRIM37 3’-UTR m6A by promoting the expression of METTL3, and the binding of m6A reader IGF2BP2 and TRIM37 3’-UTR promotes the stability of TRIM37 mRNA.

Ferroptosis, characterized by lipid peroxidation and metabolic constraints, is regulated by glutathione peroxidase (GPX4) (21). For instance, it has been reported that silencing GPX4 causes ferroptosis (22). Song et al. have reported that inhibition of GPX4 stimulated ferroptosis and enhanced breast cancer cell sensitivity to gefitinib using both in vivo and in vitro studies(23). More importantly, Ingold et al. demonstrated that catalytically inactive GPX4 confers a dominant-negative effect in male fertility (24). In this study, we found that HBs exposure significantly increased TRIM37 but decreased GPX4. Mechanism study showed that TRIM37 inhibits GPX4 expression through ubiquitination and GPX4 overexpression inhibits TRIM37’s effects on viability and ferroptosis of Sertoli cells. These results indicated a new role of TRIM37 and GPX4 in male infertility, increased our knowledge of GPX4 in ferroptosis of human Sertoli cells, and broadened our understanding of male infertility.

Ferroptosis has been involved in infections caused by HBV (25). It has been reported that chrysophanol attenuates hepatitis B virus X protein-induced hepatic stellate cell fibrosis by regulating ferroptosis (26). Neutralizing anti-HMGB1 antibodies ameliorated ferroptosis-induced inflammation (27). Inhibiting HMGB1significantly reduced ferroptosis through inhibiting oxidative stress(28). Our results indicated that HBs exposure decreased cell viability but increased ferroptosis of human Sertoli cells. The findings demonstrated novel roles of HBV infection, showing that HBs exposure increased intracellular iron concentration, promoted lipid peroxidation, decreased GSH level, and suppressed GPX4 expression, leading to decreased cell viability and increased ferroptosis of human Sertoli cells.

TRIM-containing proteins are involved in a variety of biological processes. For example, TRIM24 promotes prostate cancer cell proliferation, and aggressiveness of prostate cancer(29). More importantly, a study showed that TRIM16L, TRIM37, TRIM40, TRIM56, and TRIM59 regulate HBV viral protein expression (18), so we checked how HBs exposure affects the expression of different TRIM-containing proteins and results showed that HBs exposure significantly increased TRIM37. TRIM37 was shown to prevent the formation of centriolar protein assemblies by regulating Centrobin through ubiquitination(30). Chen et al. demonstrated that TRIM37 mediates chemoresistance of pancreatic cancer cells via ubiquitination of PTEN(31). In this study, we showed that HBs exposure dose-dependently increased TRIM37 but decreased GPX4. Mechanism study indicated that TRIM37 significantly increased the ubiquitination and degradation of GPX4, leading to an increase of ferroptosis and a decrease in the viability of human Sertoli cells. These findings indicate a critical role of TRIM37 and GPX4 in regulating ferroptosis and improving our understanding of male infertility.

HBV has been shown to involve in N6-methyladenosine modification. For example, HBV X protein interacts with METTL3 to affect m6A of viral/host RNAs(32). Elevated m6A in HCC promotes inflammation by inhibiting m6A readers (33). METTL3, as the core catalytic subunit, was reported to interact with HBx to carry out methylation activity and also modestly stimulate their nuclear import (32). In our study, we showed that HBs exposure significantly increased m6A methylation of TRIM37. Overexpressing METTL3 sharply up-regulated m6A methylation of TRIM37. Results also indicated that HBs increase the level of TRIM37 3’-UTR m6A by promoting the expression of METL3, and the binding of m6A reader IGF2BP2 and TRIM37 3’-UTR promotes the stability of TRIM37 mRNA. These results highlight the significance of METTL3/m6A modification in HBV-regulated ferroptosis of Sertoli cells and provide a new direction for the development of drugs for male infertility. Some limitations are there in this study. To further study the role of HBV/TRIM37 in male infertility, a mouse model should be employed in future research. Further studies measuring sperm quality will provide more relevant data. In conclusion, data presented here strongly suggest that HBs inhibit Sertoli cell viability through TRIM37-mediated ubiquitination of GPX4 triggering ferroptosis.

## Materials and methods

### Cell culture

Human Sertoli cells were from Guyana Biotech (Shanghai) and cultured in DMEM with 10% FBS (Invitrogen, Shanghai) at 37°C in a cell incubator containing 5% CO2.

### Exposure of Sertoli cells to HBs

HBsAg (HBs) was made in CHO cells. Secreted HBsAg was HPLC-purified. The human Sertoli cells were incubated with HBs (25, 50, and 100 mg/mL).

### Cell transfection

*TRIM37* siRNA (siR-1, 5’-GGAGAAGAUUCAGAAUGAATT-3’; siR-2,

5’-CCAGUAGUUUACUAGACAUTT-3’; siR-3,

5’-GCCUUGAUACAUGGCAGUATT-3’), *METTL3* siRNA (siR#1,

5’-GCUGCACUUCAGACGAAUUTT-3’; siR#2,

5’-GGAUACCUGCAAGUAUGUUTT-3’), and *IGF2BP2* siRNA (siR&1,

5’-CCCAGUUUGUUGGUGCCAUTT-3’; siR&2,

5’-GCGAAAGGAUGGUCAUCAUTT-3’) were synthesized by Genepharm (Shanghai, China). For ectopic expression of *TRIM37, GPX4, METTL3, or IGF2BP2*, the coding sequences were cloned into pcDNA3.1 plasmids (Addgene, USA).

Transfections were performed with Lipo2000.

### Cell viability assay

Cell viability was quantified with CCK-8 (Dojindo, Japan). Briefly, after treatment, CCK-8 was added and incubated for 1h. The absorbance at 450nm was determined using a microplate reader.

### Measurement of glutathione (GSH), MDA, and iron content

GSH and MDA were measured with Reduced Glutathione Assay Kit (A006-1) and Malondialdehyde Assay Kit (A003-1-2) (Nanjing Jiancheng Bio.). The Iron Assay Kit (Abcam; ab83366) was used for the detection of iron content according to the instructions of the manufacturer.

### Lipid peroxidation assessment using C11-BODIPY

The cell suspension was added with 10 mM C11-BODIPY (Thermo Fisher Scientific, USA) and incubated for 30 min at 37°C in the dark. After two-time washing with PBS, the fluorescence intensity of the samples was measured by Accuri™ C6 flow cytometer (BD Biosciences).

### Quantitative RT-PCR

RNAs were collected from the human Sertoli cells using TRIzol (Life Technologies, Waltham, MA) and cDNA was synthesized. qPCR with SYBR green PCR master mix (ABI, Foster, CA) was performed in an ABI 9700 real-time PCR system. The primers used are as follows (5’-3’): TRIM16L-F: GCCGAGATGGAGAAGAGTAAG, TRIM16L-R: GCTGAACAATGGCAGACAC; TRIM37-F: TGGACTTACTCGCAAATG, TRIM37-R: ATCTGGTGGTGACAAATC; TRIM40-F: ATGCCCTCAGCCACTACAAG, TRIM40-R: TCCCGTGGTCTACCTGAAAC; TRIM56-F: GAAACGCTTCTCCCTCAAC, TRIM56-R: CTTGCCTCCAGGAATGAAC; TRIM59-F: TGCCTTACCATAGGTCAAC, TRIM59-R: GATTGCCAACATCACAGAG; METTL3-F: CCTTTGCCAGTTCGTTAGTC, METTL3-R: TCCTCCTTGGTTCCATAGTC; IGF2BP2-F: CGGGAGCAAACCAAAGACC, IGF2BP2-R: GCAAACCTGGCTGACCTTC; GPX4-F: AACCCAAGGGCAAGGGCATC, GPX4-R: ACCACGCAGCCGTTCTTGTC; b-actin-F: TGGCATTGCCGACAGG, b-actin-R: GCATTTGCGGTGGACG. Gene fold changes were determined by 2−ΔΔCT.

### Western blot analysis

Proteins were separated on SDS-PAGE, immunoblotted to membranes, blocked, and incubated with primary antibodies against TRIM37 (Abcam; ab264190), GPX4 (Abcam; ab125066), METTL3 (Abcam; ab195352), IGF2BP2 (Abcam; ab129071), or b-actin (Cell Signaling Technology; #4970) at 4°C overnight, followed by goat anti-rabbit IgG (Beyotime, China; A0208) antibody for 1 h at room temperature. Western blotting band intensity was quantified by Quantity One software.

### Co-immunoprecipitation assay

Cell lysates were probed with anti-TRIM37 (Abcam; ab264189), anti-GPX4 (Proteintech; 14432-1-AP), or normal IgG at 4°C for 12h, followed by 2 h Protein A/G Agarose beads at 4°C for 2h. Immunocomplexes were washed and immunoblotted with anti-TRIM37 (Abcam; ab264190), anti-GPX4 (Abcam; ab1250663), and anti-ubiquitin (ab7780; Abcam) antibodies.

### m6A level analysis

The total levels of m6A were examined with the m6A RNA Methylation Assay Kit (Abcam, MA, USA).

### RNA immunoprecipitation (RIP) assay

RIP assay was done with Magna RIP RNA-Binding Protein Immunoprecipitation Kit (Millipore, USA). Quantitative RT-PCR was used for determining total RNA levels.

### Luciferase reporter assay

The 3¢-UTR of TRIM37 containing the wild-type m6A motifs as well as mutant m6A motifs (m6A was replaced by T) was synthesized by General Tech. (Shanghai), and ligated into the pGL3-basic vector (Promega). Then, wild-type or mutant pGL3-TRIM37-3¢-UTR and URL-TK renilla vectors were co-transfected into Sertoli cells overexpressing METTL3. After 48 h, dual-luciferase activities were measured by the Dual-Luciferase Assay System (Promega, USA).

## Data analysis

Data analysis was performed with GraphPad Prism 8.0.2. All data were presented in mean ± SD. Significances between the 2 groups were determined by unpaired Student’s t-tests, and one-way or two-way ANOVA was performed to compare among 3 or more groups. p < 0.05 was considered statistically significant.

